# CAMSAP3 Forms Dimers via its α-helix Domain that Directly Stabilize Non-centrosomal Microtubule Minus Ends

**DOI:** 10.1101/2024.05.31.596772

**Authors:** Yuejia Li, Rui Zhang, Jinqi Ren, Wei Chen, Zhengrong Zhou, Honglin Xu, Dong Li, Haisu Cheng, Qi Xie, Wei Ji, Wei Feng, Xin Liang, Wenxiang Meng

## Abstract

Microtubules are vital components of the cytoskeleton. Their plus ends are dynamic and respond to changes in cell morphology, while the minus ends are stable and serve a crucial role in microtubule seeding and maintaining spatial organization. In mammalian cells, the calmodulin-regulated spectrin-associated proteins (CAMSAPs), play a key role in directly regulating the dynamics of non-centrosomal microtubules minus ends. However, the molecular mechanisms are not yet fully understood. Our study reveals that CAMSAP3 forms dimers through its C-terminal α-helix; this dimerization not only enhances the microtubule-binding affinity of the CKK domain but also enables the CKK domain to regulate the dynamics of microtubules. Furthermore, CAMSAP3 also specializes in decorating at the minus end of microtubules through the combined action of the microtubule-binding domain (MBD) and the C-terminal α-helix, thereby achieving dynamic regulation of the minus ends of microtubules. These findings are crucial for advancing our understanding and treatment of diseases associated with non-centrosomal microtubules.

**Significance:** Our study reveals the molecular mechanism of how CAMSAP3, a key regulator of non-centrosomal microtubule dynamics, directly regulates the dynamics of non-centrosomal microtubule minus ends through CKK domain. CAMSAP3 forms dimers through its C-terminal α-helix, which enhances the CKK domain of CAMSAP3 binding to microtubule minus ends and confers stability of them. This finding is not limited to CAMSAP3, but can also be applied to the understanding of the regulation of non-centrosomal microtubule minus end stability by CAMSAP family proteins. Our findings deepen our comprehension of cellular structure and function, offering insights into the role of microtubules in cellular integrity and disease. This study fills a significant knowledge gap and lays the foundation for future research into the complex balance of microtubule dynamics required for cellular health and disease prevention.

## Introduction

Microtubules are a crucial component of the cytoskeleton, and they play a vital role in various biological processes such as cell movement and establishment of polarity (1). The plus ends of microtubules are highly dynamic, while the minus ends are relatively stable (2, 3). These stable minus ends can act as a foundation for microtubule regeneration, which helps maintain the organization and effectiveness of the microtubule networks (4–7). The stability of microtubule minus ends is highly regulated by its binding proteins (8). Currently, it is known that the γ-tubulin ring complex (γ-TuRC) binds α-tubulin at the minus end of centrosomal microtubules, providing a basis for maintaining microtubule stability (9–12). Current studies have shown that proteins like the CAMSAP family, ASPM, CEP170, and KANSL complex can recognize and regulate the minus end dynamics of non-centrosomal microtubules (13–17). However, the molecular mechanisms underlying their dynamic regulation are not well understood.

The CAMSAP protein family, essential for anchoring and controlling the dynamics at the minus ends of microtubules, is conserved across mammals and invertebrates, and include PTRN-1 and Patronin (18–20). These proteins are structurally characterized by an N-terminal CH domain, three CC domains, and a C-terminal CKK domain (21). The previous study, incorporating both in vivo and in vitro studies, has identified the CKK domain as pivotal for CAMSAP proteins’ capacity to recognize microtubule minus ends. This domain is known to specifically bind and adhere to interprotofilament tubulin junctions within microtubules, prompting their conformational changes (22, 23). The CKK domain plays a prerequisite role in the localization of CAMSAP family proteins at the minus end of microtubules and in regulating their stability (16, 22–24). However, the current understanding does not fully support the function of CAMSAP family proteins, because it cannot individually modulate microtubule dynamics, and also cannot distribute to the minus ends of microtubules like the full length of CAMSAP2 or CAMSAP3 (16). Further complexity arises when comparing within the CAMSAP family: while the CKK domain’s presence is highly conserved across CAMSAP1-3, both CAMSAP2 (KIAA1708) and CAMSAP3 (Nezha, KIAA1543) exhibit significantly enhanced proficiency in limiting minus end minus end dynamics as compared to CAMSAP1 (15, 16). These indicate that the mechanism by which CAMSAP proteins modulate dynamics of the non-centrosomal microtubule minus end is intricate and not yet fully understood.

Our study advances the understanding of how CAMSAP3 dynamically regulates and specifically localizes at microtubule minus ends, highlighting the critical role of the α-helix (980^th^-1005^th^ a.a.) domain adjacent to the CKK domain. We have found that this α-helix facilitates the dimerization of CAMSAP3, which not only strengthens the CKK domain’s microtubule affinity but also equips it with the ability to modulate minus end dynamics. Furthermore, our data reveal that this α-helix, in conjunction with the microtubule binding domain (MBD), markedly contributes to the precise decoration of microtubule minus ends by CAMSAP3. The synergy between the localization to microtubule minus end specificity and the stabilizing effect of CKK dimerization empowers CAMSAP3 to regulate these ends dynamically. Our study also suggests that differences between the α-helical domains significantly impact the heterogeneity of CAMSAP family proteins. The insights we gained contribute to understanding the regulation of non-centrosomal microtubules.

## Results

### Specific Regulation of Microtubule Minus End Dynamics by CAMSAP3 and Its Characteristic Localization

To further study the molecular mechanism by which CAMSAP family proteins regulate microtubule minus end dynamics, we first established an experimental system for microtubule stabilization in vitro (Figure 1 A). We purified GFP-CAMSAP3 and GFP-CKK and conducted in vitro microtubule dynamics experiments (Figure 1 B-F). Results showed that CAMSAP3 decorates minus ends of microtubules and significantly modulates their dynamics (Figure 1 E-H). In contrast, GFP-CKK was bound to the minus ends and lattice of microtubules but cannot regulate microtubule dynamics like full-length CAMSAP3 (Figure 1 F-H). Consistent results were obtained with cultured cells overexpressing full-length CAMSAP3 or just its CKK domain (Supplemental Figure 1 A and B), which aligns with previous reports (15, 16, 22).

**Figure 1.**
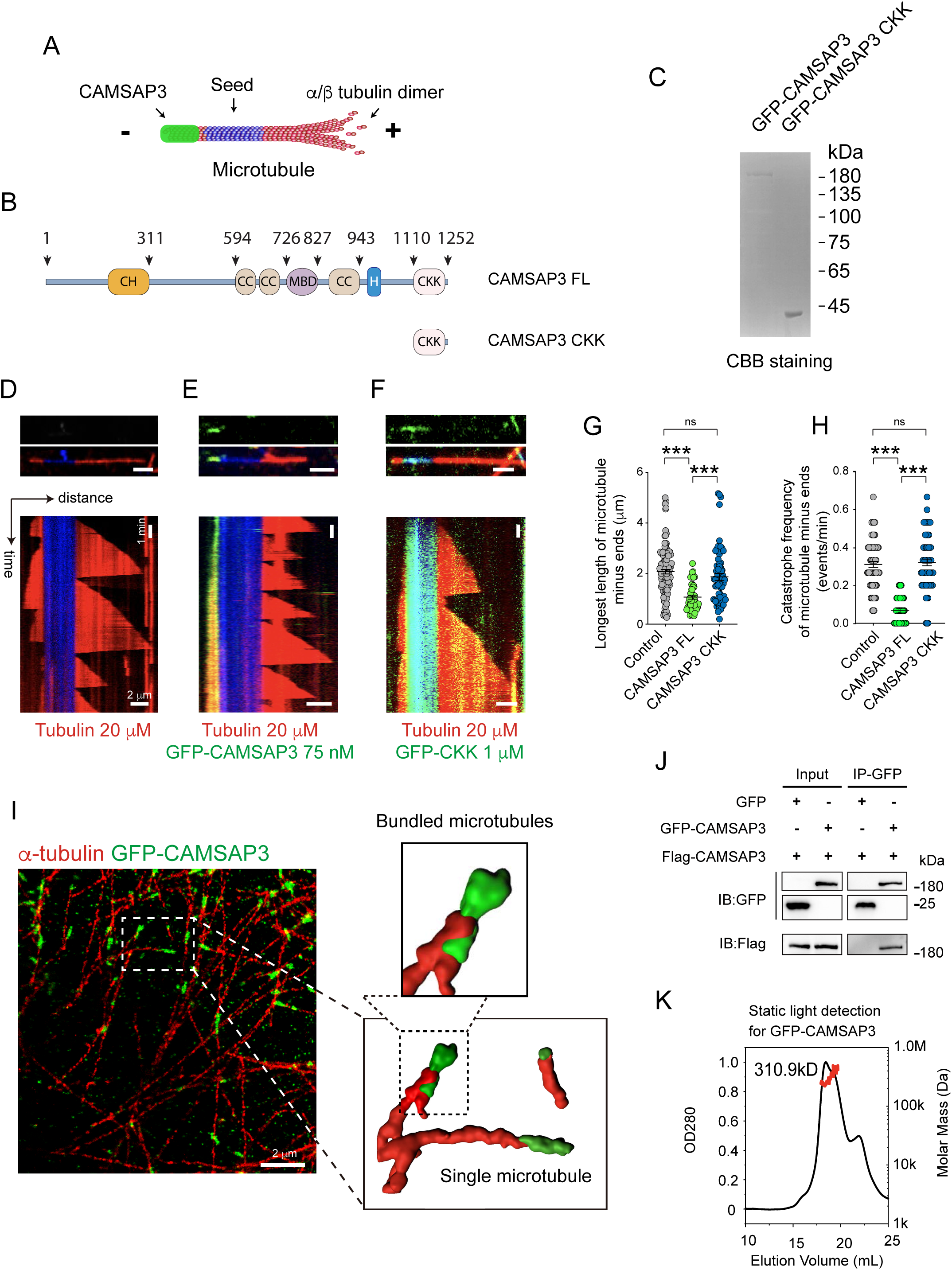
CAMSAP3 decorates at microtubule minus ends and regulates their dynamics. **(A)** Scheme of the microtubule minus ends dynamic assay. GMPCPP microtubule seeds were attached to an Avidin-coated coverslip. Tubulin was allowed to polymerize in the presence of CAMSAP3 proteins. **(B)** Schemes of the domain organization of CAMSAP3 full-length and CAMSAP3 CKK. **(C)** Coomassie Brilliant Blue staining of a gel with GFP-CAMSAP3 and GFP-CAMSAP3 CKK purified from Sf9 cells. **(D-F)** Representative kymographs of control (D), GFP-CAMSAP3 (E), and GFP-CKK (F) decorated microtubules. Scale bars: horizontal, 2 μm; vertical, 1 min. **(G)** Quantification of the longest length of microtubule minus ends. n = 112 microtubules, control; n = 48 microtubules, GFP-CAMSAP3; n = 66 microtubules, GFP-CKK. **(H)** Quantification of the catastrophe frequency of microtubule minus ends decorated with CAMSAP3 proteins. n = 68 microtubules, control; n = 46 microtubules, GFP-CAMSAP3; n = 67 microtubules, GFP-CKK. Data represent mean ± SEM. **(I)** A representative ROSE-Z super-resolution microscope image of microtubules in Caco-2 cells. α-tubulin is shown in red and GFP-CAMSAP3 is displayed in green. A reconstructed image of CAMSAP3 exhibited a decorating pattern at the microtubule minus ends visualized in the right plane. Scale bar: 2 μm. **(J)** Co-immunoprecipitation assays were performed with the extracts of HEK293T cells co-expressing GFP-CAMSAP3 and Flag-CAMSAP3 to confirm the self-binding of CAMSAP3. **(K)** Static light detection of GFP-CAMSAP3. The primary peak value corresponds to the dimer form of GFP-CAMSAP3.

Next, we observed the distribution of GFP-CAMSAP3 in HeLa cells using the ROSE-Z super-resolution microscope. The results revealed that CAMSAP3 decorates and wraps around one end of microtubules (Figure 1 I). Notably, it could bundle microtubules and wrap around more than one microtubule (the enlarged portion of Figure 1 I). We also observed a similar structure in cells that overexpress CAMSAP2, but it was difficult to identify it in cells overexpressing CAMSAP1, which seems to prefer localizing to the lattice of microtubules (Supplemental Figure 1 C and D). Previous studies have also reported the distinct localization characteristics of CAMSAP2 and CAMSAP3 compared to CAMSAP1 (15, 16, 22). Further analysis of the mechanism underlying the differential localization tendencies of CAMSAPs is necessary to understand its molecular regulatory mechanism on microtubules.

### CAMSAP3 Predominantly Exists as a Dimer

CAMSAP3 could decorate and wrap around the minus ends of microtubules (Figure 1 I), and it also can induce microtubule bundling (7, 25), inspiring us that CAMSAP3 has the potential to cross-link microtubules through self-interactions. To verify this possibility, immunoprecipitation assays using different tagged versions of CAMSAP3 confirmed that CAMSAP3 can self-bind (Figure 1 J). Considering the limitation of immunoprecipitation assays, which make it difficult to distinguish between homodimerization and coimmunoprecipitation via a common other scaffolding protein, we used purified GFP-CAMSAP3 (Figure 1 C), performed static light detection, and identified a primary peak at 310.9 kD (Figure 1 K). Given that GFP-CAMSAP3’s molecular weight is approximately 162 kD, this peak corresponds to the dimer. Similarly, the same experiments demonstrated that CAMSAP1 or CAMSAP2 also exhibit self-association capabilities (Supplemental Figure 1 E and F). Therefore, these data suggest that CAMSAPs could self-associate and that this property is conserved within the protein family.

To investigate the self-interactions of CAMSAP3, we generated and transfected a series of mutants into HEK293T cells (Figure 2 A and Supplemental Figure 2 A). We examined their interactions using co-immunoprecipitation experiments and showed that truncated mutants containing their N- or C-termini can interact individually (Figure 2 B and Supplemental Figure 2 B). However, the interaction of their C-terminus requires the simultaneous presence of α-helix and CKK domain (Supplemental Figure 2 C). Interestingly, the characteristics of these interactions are highly conserved within the CAMSAP family (Supplemental Figure 3). While the interaction domains via their N-termini are relatively conserved for CAMSAP family proteins (Supplemental Figure 3 A, B, E, and F), their C-terminal interactions occur through different domains (Supplemental Figure 3 C, D, G, and H). These data suggest that while self-interaction is a common feature among CAMSAP family proteins, each protein has its own unique characteristics.

**Figure 2.**
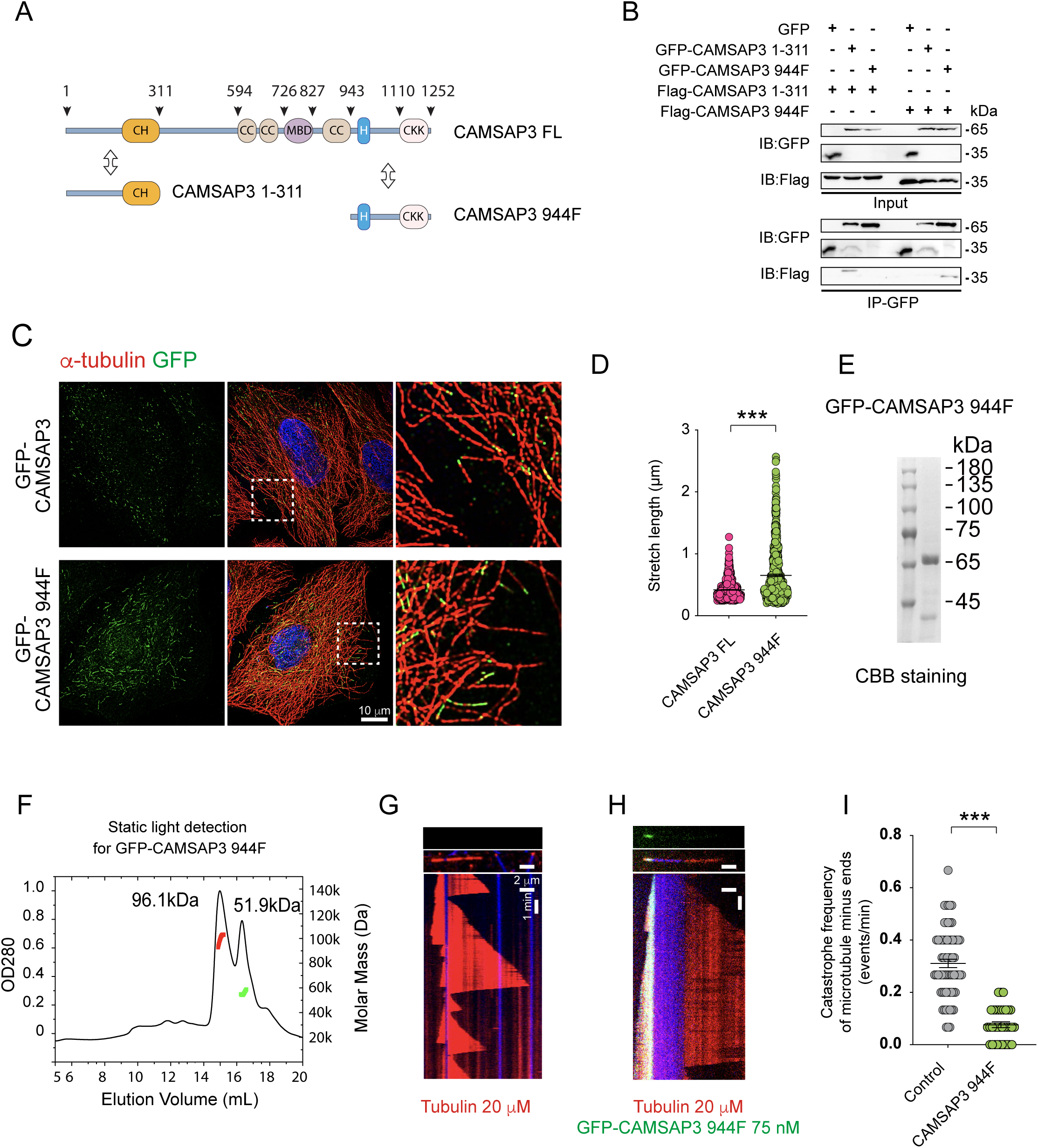
C-terminal dimerization of CAMSAP3 is sufficient for its microtubule minus ends localization and dynamic regulation. **(A)** Schemes of the domain organization of CAMSAP3 full-length and truncated mutants were used to determine the dimerization region of the CAMSAP3. **(B)** Co-immunoprecipitation assays were performed to confirm the self-binding of CAMSAP3 N- or C-terminal. **(C)** Immunostaining for α-tubulin (red) and GFP (green) in GFP-CAMSAP3 and GFP-CAMSAP3 944F transfected HeLa cells. Scale bar: 10 μm. **(D)** Quantification of GFP-CAMSAP3 and GFP-CAMSAP3 944F signal length in HeLa cells. n = 683 from three cells, GFP-CAMSAP3; n = 554 from three cells, GFP-CAMSAP3 944F. Data represent mean ± SEM. **(E)** Coomassie Brilliant Blue staining of a gel with GFP-CAMSAP3 944F protein purified from Sf9 cells. **(F)** Static light detection of GFP-CAMSAP3 994F protein. The primary peak value corresponds to the dimer form of GFP-CAMSAP3 944F. **(G and H)** Representative kymographs of microtubule dynamics of control (G) or in the presence of GFP-CAMSAP3 944F (H). Scale bars: horizontal, 2 μm; vertical, 1 min. **(I)** Quantification of microtubule minus ends catastrophe frequency for the experiments shown in (G and H) n = 68 microtubules, control; n = 34 microtubules, GFP-CAMSAP3 944F. Data represent mean ± SEM.

To understand the functions of the N-terminus and C-terminus of CAMSAP3, we conducted an in vitro microtubule dynamics experiment using a purified truncated mutant of the proteins (Supplemental Figure 4 A). Results showed that the lack of the N-terminus did not impact the distribution or dynamic regulation of microtubule minus ends by CAMSAP3 (Supplemental Figure 4 B-F). Notably, the C-terminus (595F or 944F) still decorates the minus ends of microtubules (Supplemental Figure 4 G and H, Figure 2 C and D). Static light detection using purified protein showed that the primary peak at 96.1 kD and 51.9 kD corresponds to the dimer and monomer form of GFP-CAMSAP3 944F (Figure 2 E and F). Additionally, it also can decrease the catastrophe rate of the minus end of microtubules (Figure 2 G-I).

All these results suggest that the C-terminus of CAMSAP3 is necessary and sufficient for regulating microtubule minus end dynamics and ensuring its decorating on the minus end of microtubules.

### The α-helix Domain Mediates C-terminal Dimerization of CAMSAP3

We created a C-terminal truncation mutant of CAMSAP3 to investigate its function in more detail. To avoid interference with N-terminal interaction, we also deleted its N-terminus and α-helical region (CAMSAP3 Δ1-311 & 944-1110) (Supplemental Figure 5 A). We found this deletion mutant of CAMSAP3 lost its ability to bind to microtubules (Supplemental Figure 5 D-E). This result is consistent with the microtubule-binding region D2S of CAMSAP3 (976^th^ to1025^th^ a.a.) reported by Atherton et.al, as well as the D2 region of CAMSAP3 (974^th^ to 1111^st^ a.a.), which is responsible for microtubule binding and dynamics regulation (22, 26). Furthermore, the mutant impaired its ability to interact with itself (Supplemental Figure 5 B-C). This suggests that the C-terminus amino acid sequence from 944^th^ to 1110^th^ a.a. of CAMSAP3, is essential not only for its binding to microtubules but also for its ability to engage in self-interaction.

To further study the molecular mechanism of dimer formation, we used AlphaFold2 to predict the sequences involved in interactions within a specific region (944^th^ to 1110^th^ a.a. of CAMSAP3, Figure 3 A and B) (27). Our results suggested that the α-helix domain (985^th^ -1001^st^ a.a.) in the C-terminus of CAMSAP3 has strong hydrophobic interactions and forms a reverse dimer (Figure 3 B). It is worth noting that the arrangement of leucine in this area is consistent with the leucine zipper structure of the α-helix (Figure 3 C), which forms a dimer (28). We then mutated the leucines at positions 993^rd^ and 997^th^ a.a. and found that these mutants have a significantly reduced propensity to interact by immunoprecipitation (Figure 3 D).

**Figure 3.**
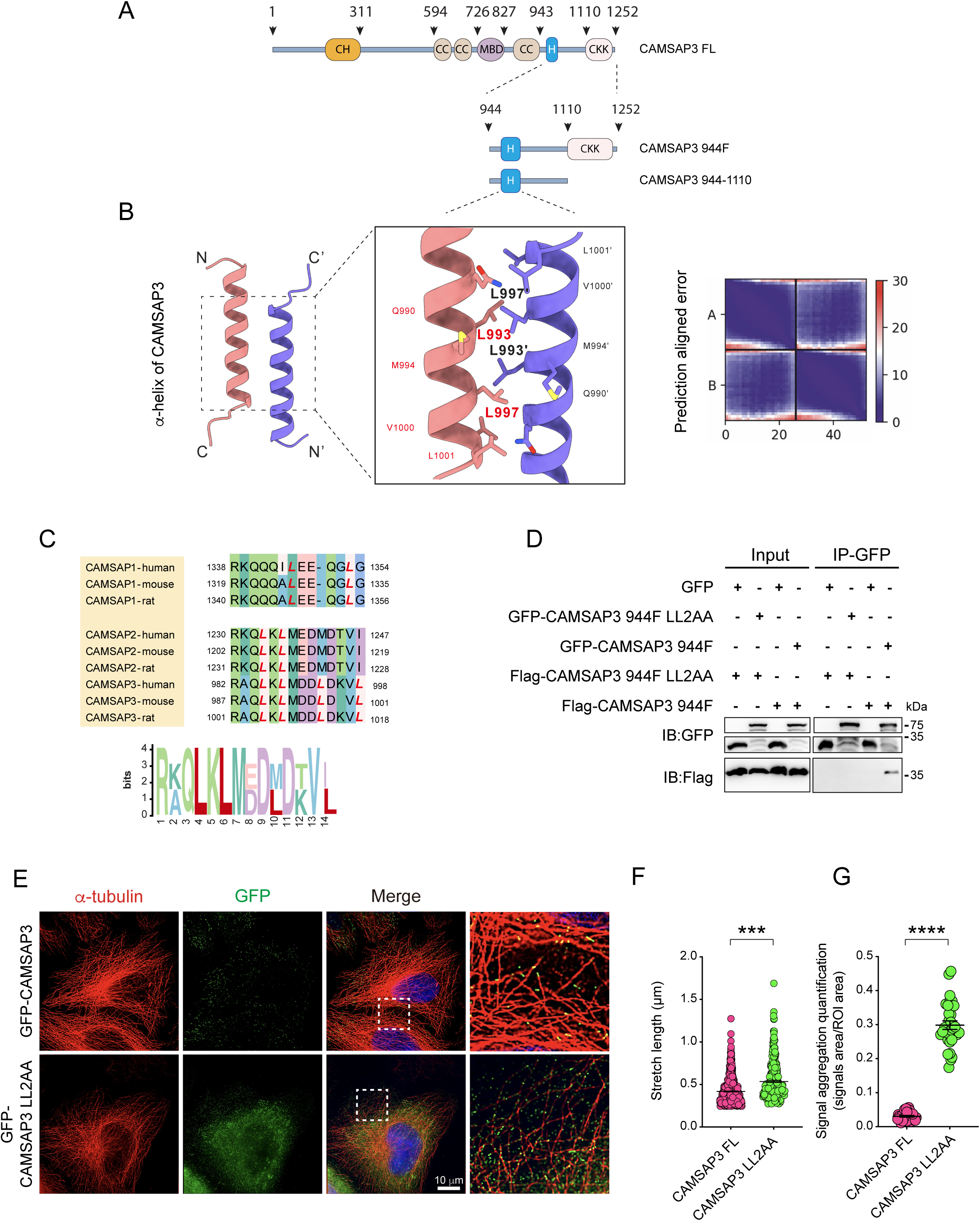
CAMSAP3 forms dimers through the *α*-helix at its C-terminus. **(A)** Schemes of the domain organization of CAMSAP3 full-length and truncated mutants mediate the dimerization of the CAMSAP3 C-terminal. **(B)** Dimerization structure of CAMSAP3 α-helix predicted by AlphaFold2. **(C)** Alignment of amino acid sequences of α-helix in the C-terminal of CAMSAP family proteins. **(D)** Co-immunoprecipitation assays were performed to detect the self-binding of CAMSAP3 944F and CAMSAP3 944F LL2AA. **(E)** Immunostaining for α-tubulin (red) and GFP (green) in GFP-CAMSAP3 and GFP-CAMSAP3 LL2AA transfected HeLa cells. **(F)** Signal length of GFP-CAMSAP3 and GFP-CAMSAP3 LL2AA was quantified in HeLa cells. n = 683 from three cells, CAMSAP3 FL; n = 303 from three cells, CAMSAP3 LL2AA. Data represent mean ± SEM. **(G)** Signal aggregation quantification of GFP-CAMSAP3 and GFP-CAMSAP3 LL2AA. Three regions of interest avoid the nucleus were selected in one cell. For each ROI, the value represents the ratio of the GFP signal area to the whole ROI area. n = 30 from 10 cells, both of CAMSAP3 FL and CAMSAP3 LL2AA. Data represent mean ± SEM.

Next, we mutated these sites in full length of CAMSAP3 and transfected it into HeLa cells, examined it by immunostaining, and results showed that the portion of microtubule minus ends decorated by CAMSAP3 in cells is significantly reduced, and CAMSAP3 is evenly distributed in the cytoplasm or along microtubules, suggesting that the binding of microtubules by CAMSAP3 is closely related to the α-helix domain (Figure 3 E-F). Interestingly, the leucine arrangement characteristics of the α-helical domain are not conserved among CAMSAP family proteins. CAMSAP2 and CAMSAP3 have leucine zipper deconstruction, but CAMSAP1 does not (Figure 3 C).

### Dimerization of CKK Can Improve its Ability to Bind Microtubules

Proteins’ dimerization is crucial for their functionality and activity, especially in microtubule-binding proteins like MAP2 and APC (29–31). To investigate the implications of CKK domain dimerization (Figure 4 A), we utilize the FKBP-FRB system to create artificial dimers that can be induced to form stable dimers using rapamycin (32).

**Figure 4.**
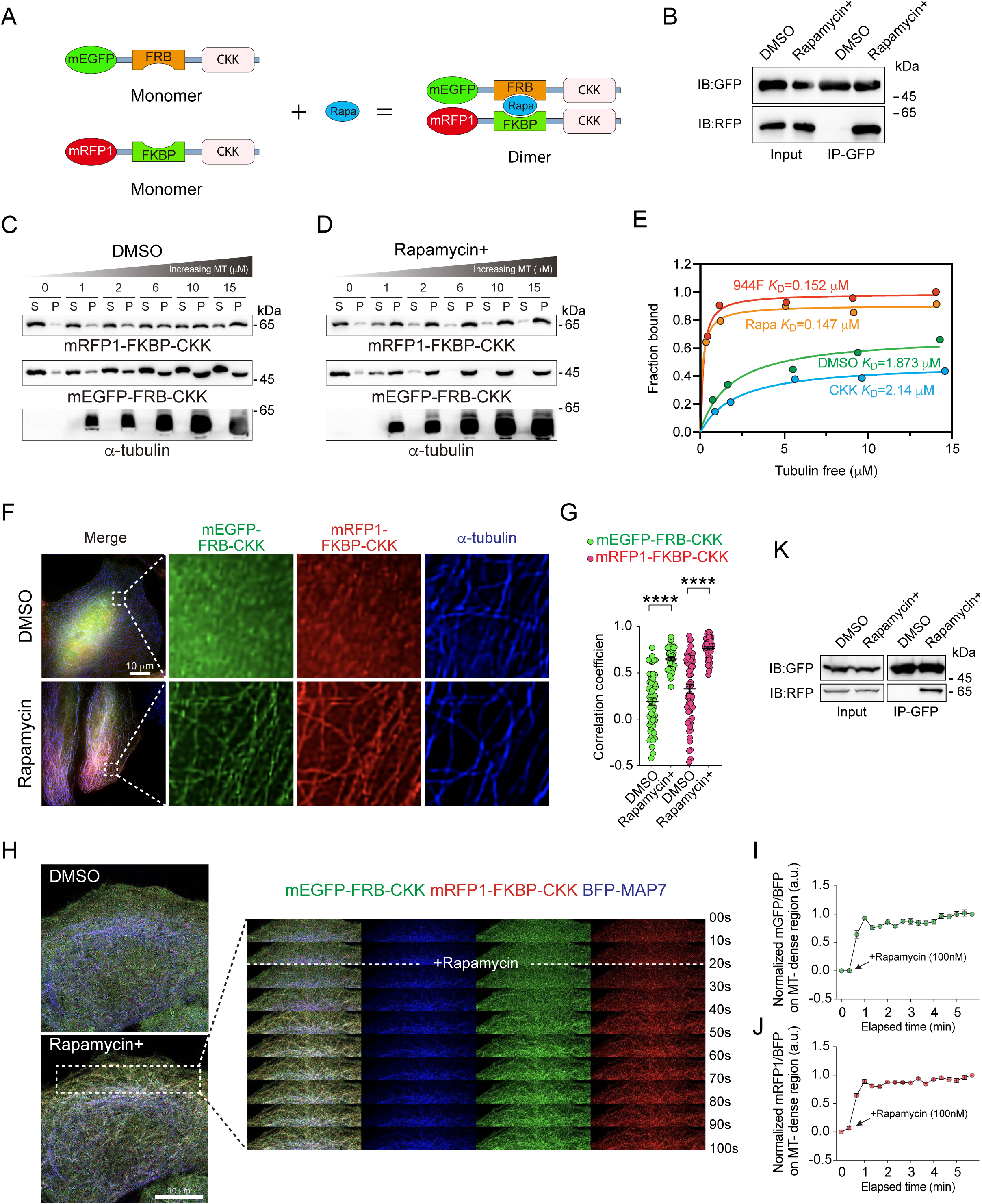
The dimerization of CKK can significantly enhance its binding affinity to microtubules. **(A)** Schemes of the FRB-FKBP dimerization system used to construct artificial CKK dimers. **(B)** Pull-down assays were performed to confirm the dimerization of purified CKK proteins. **(C and D)** Microtubule co-sedimentation assays were performed to detect the binding affinity of CKK monomer (C) and dimer (D) to microtubules. Taxol-stabilized microtubules were assembled in vitro. Supernatant and pellet were analyzed by western blotting with the indicated antibodies. **(E)** The dissociation constant (*K*_D_) of CKK monomer, dimer, control, and 944F were calculated from microtubule co-sedimentation assays. **(F)** Immunostaining for mEGFP-FRB-CKK (green), mRFP1-FKBP-CKK (red), and α-tubulin (blue) in DMSO and 100 nM rapamycin-treated HeLa cells. Scale bar: 10 μm. **(G)** Co-localization of CKK and microtubules in DMSO and 100 nM rapamycin-treated HeLa cells analyzed from experiments shown in (F), n = 54 cells, DMSO; n = 44 cells, rapamycin. Data represent mean ± SEM. **(H)** Live cell images of mEGFP-FRB-CKK (green), mRFP1-FKBP-CKK (red), and BFP-MAP7 (blue) of HeLa cells at indicated time points after 100 nM rapamycin was added. **(I and J)** Quantification of the mEGFP-FKBP-CKK (I) or mRFP1-FRB-CKK (J) translocation to BFP-MAP7 in 4H. Two regions of interest at microtubule dense regions were selected. For each ROI, the last YFP/CFP is normalized to 1 and the initial YFP/CFP is normalized to 0. n = 12 from 6 cells. Data represent mean ± SEM. **(K)** Co-immunoprecipitation assays were performed to detect the dimerization of CKK in DMSO or 100 nM rapamycin-treated cells.

First, we purified these proteins and verified their interaction (Supplemental Figure 6 A and Figure 4 B). Notably, our microtubule co-sedimentation assay revealed that dimerization of CKK significantly enhances its binding capacity to microtubules (Figure 4 C and D). The maximum binding increased from 0.692 to 0.904, while the dissociation coefficient decreased from 1.873 µM to 0.147 µM (Figure 4 E). This indicates that the dimerization has improved CKK’s binding affinity with microtubules by almost 13-fold. We also performed experiments to analyze the binding of microtubules with CAMSAP3-C terminus (944F) and CKK (Supplemental Figure 6 B). The result showed that the maximum binding and dissociation coefficient of CAMSAP3-C terminus (944F) with microtubules were 0.986 and 0.152 µM, respectively (Supplemental Figure 6 C and D, Figure 4 E). These values are consistent with those of the CKK dimer form, indicating that the dimerization of CKK is comparable to the actual affinity of CAMSAP3-C terminus (944F) to microtubules.

To ensure that our findings were not limited to in vitro conditions, we co-transfected mRFP1-FKBP-CKK and mEGFP-FRB-CKK into cells (Figure 4 F-J). These results showed that when induced with rapamycin, the colocalization of mRFP1-FKBP-CKK or mEGFP-FRB-CKK with microtubules was significantly improved. This was confirmed through immunostaining (Figure 4 F and G), and time-lapse microscopy observations (Figure 4 H-J). Furthermore, these results were also confirmed by immunoprecipitation experiments (Figure 4 K).

All these results strongly suggest that dimerization of the CKK domain promotes its binding to microtubules.

### Dimerized CKK Can Reduce the Frequency of Catastrophe of Microtubules

CAMSAP3’s regulation of microtubule minus ends dynamics is linked to its C-terminus (15, 16), which comprises an α-helix and CKK domain (Figure 2 C, I and J). Additionally, our results suggest that the α-helix may facilitate the dimerization of the C-terminus of CAMSAP3 (Figure 3). Therefore, it is highly implied that the dimerization of CKK plays a crucial role in the dynamic regulation of microtubule minus ends by CAMSAP3.

To test this idea, we used purified mRFP1-FKBP-CKK and mEGFP-FRB-CKK to study the dynamic regulation of microtubule minus ends in monomers and dimers (Supplemental Figure 6 A and Figure 5 A). It is noted that the individual CKK domain cannot decorate the minus ends of microtubules (16, 22). Therefore, regardless of whether it is the CKK monomer or dimer, both can bind to the microtubule lattice as protein concentrations increase. We found that both the minus and plus ends of microtubules exhibited a higher frequency of catastrophe in the monomer case, whereas this occurred less frequently in rapamycin-induced dimers (Figure 5 B and C). This result demonstrates that the dimer of CKK is indeed related to the dynamic regulation of microtubules by CAMSAP3.

**Figure 5.**
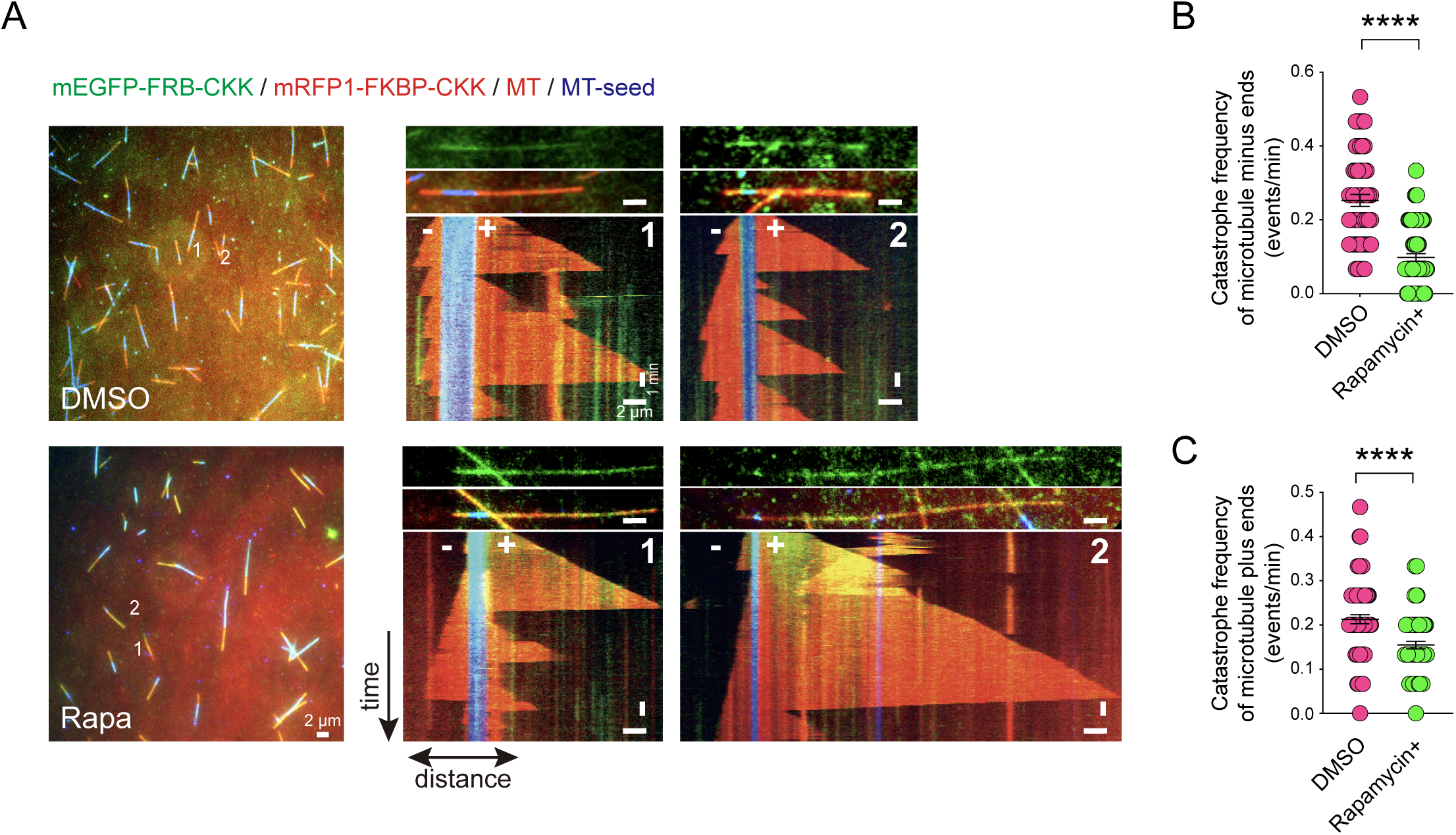
The dimerization of CKK can stabilize microtubule plus ends and minus ends. **(A)** Representative kymographs of microtubule dynamics of CKK monomer and CKK dimer decorated microtubules. Scale bars: horizontal, 2 μm; vertical, 1 min. **(B)** Quantification of microtubule minus ends catastrophe frequency for the experiments shown in (A) n = 56 microtubules, DMSO; n = 72 microtubules, 100 nM rapamycin. Data represent mean ± SEM. **(C)** Quantification of microtubule plus ends catastrophe frequency for the experiments shown in (A) n = 73 microtubules, DMSO; n = 69 microtubules, 100 nM rapamycin. Data represent mean ± SEM.

To confirm our conclusion, we also used GCN4, a tag for forming artificial dimers (33), fused with CKK (Supplemental Figure 6 E). We transfected GFP-GCN4-CKK and control constructs into HeLa cells and immunostained them with anti-α-tubulin and EB1 antibodies (Supplemental Figure 6 F). Compared with CKK, the microtubule-binding ability of GCN4-CKK was significantly increased (Supplemental Figure 6 G). It should be noted that in cells overexpressing GCN4-CKK, the number of EB1 was significantly decreased compared to cells overexpressing GFP-CKK and GFP-GCN4 (Supplemental Figure 6 H). EB1 is the marker for growing microtubule plus ends, and its reduced quantity suggests a more stable state for microtubules. All these data suggest dimerization confers the ability of CKK to regulate microtubule dynamics.

### The α-helix Regulates CAMSAP3 Decorating at the Minus Ends of Microtubules

In the above experiment, it was observed that the plus ends of microtubules also exhibited a phenotype of reduced collapse frequency (Figure 5 C). It is speculated that regardless of whether it is in vivo or in vitro, CKK tends to bind to the lattice of microtubules, including the plus ends of microtubules (Figure 5 A, 4 F and H, and Supplemental Figure 1 B). This suggests that dimerized CKK requires assistance to locate the minus ends of microtubules for specific regulation.

To investigate this matter further, we introduced a CAMSAP3 truncation mutant starting from the N-terminus into HeLa cells and used immunostaining to observe its localization at microtubule minus ends (Supplemental Figure 7 A). Our findings indicate that mutants truncated up to the first coiled-coil domain (595F) still retain the ability to decorate at microtubule minus ends (Supplemental Figure 4 G). However, mutants starting from the MBD domain (727F) exhibit a weakened ability to decorate at minus ends, although this ability persists to some extent in the mutant truncated to the 980^th^ a.a. (980F). The mutant truncated at amino acid 985 (985F) completely loses this ability, showing a weak, diffuse microtubule-binding phenotype like CKK (Supplemental Figure 1 B). These observations suggest that multiple factors influence the localization of CAMSAP3 at microtubule minus ends. It is important to note that the amino acid sequence between 980^th^ and 984^th^ a.a. plays a crucial role. Even with the MBD and CKK domains, the depletion mutation CAMSAP3-Δ980-984 cannot decorate at the microtubule minus end and loses its ability to bind to microtubules (Supplemental Figure 7 B). In addition, these mutants still retain their self-interacting properties (Supplemental Figure 8). These data indicate that α-helix is also necessary for CAMSAP3 decorating at the microtubule’s minus end.

### α-helix is important for Heterogeneity of CAMSAP Family Proteins

Our research results suggest that α-helix is very important for the localization of CAMSAP3 at the minus end of microtubules (Supplemental Figure 7 B and C, Figure 6 A and B), and correspondingly, the C-terminus of CAMSAP1 and CAMSAP2 appear to be evenly distributed along microtubules and cannot specifically localize to the minus ends of microtubules (Figure 6 C-E). This result suggests that the different amino acid characteristics of the corresponding domain of α-helix may be an important reason for determining the differences in the distribution patterns of the CAMSAP family on the minus ends of microtubules.

**Figure 6.**
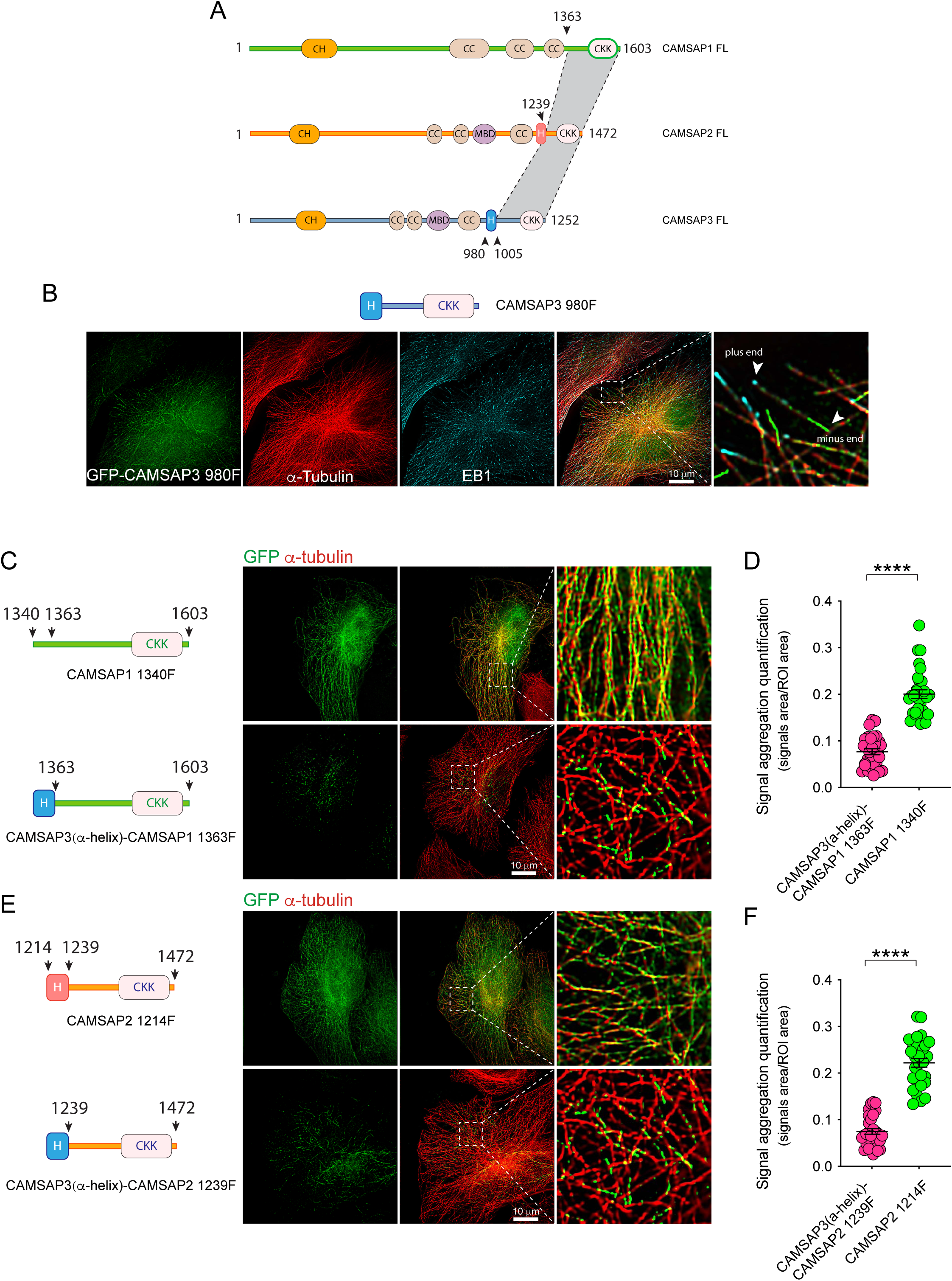
The *α*-helix of CAMSAP3 enable CAMSAPs to localize at microtubule minus ends. **(A)** Schemes of the conserved regions (in gray) in CAMSAPs that are adjacent to the α-helix in C-termini. **(B)** Immunostaining for α-tubulin (red), GFP (green), and EB1 (blue) in GFP-CAMSAP3 980F transfected HeLa cells. GFP-CAMSAP3 980F decorates microtubule minus ends. **(C)** Immunostaining for α-tubulin (red) and GFP (green) in GFP-CAMSAP1 1340F and GFP-CAMSAP3(α-helix)-CAMSAP1 1363F transfected HeLa cells. The α-helix of CAMSAP3 enables the CAMSAP1 C-terminal to localize at microtubule minus ends. **(D)** Signal aggregation quantification of GFP-CAMSAP1 1340F and GFP-CAMSAP3(α-helix)-CAMSAP1 1363F in HeLa cells. n = 30 from 10 cells, both of CAMSAP1 1340F and CAMSAP3(α-helix)-CAMSAP1 1363F. Data represent mean ± SEM. **(E)** Immunostaining for α-tubulin (red) and GFP (green) in GFP-CAMSAP2 1214F and GFP-CAMSAP3(α-helix)-CAMSAP2 1239F transfected HeLa cells. The α-helix of CAMSAP3 enables the CAMSAP2 C-terminal to localize at microtubule minus ends. **(F)** Signal aggregation quantification of GFP-CAMSAP2 1214F and GFP-CAMSAP3(α-helix)-CAMSAP2 1239F in HeLa cells. n = 30 from 10 cells, both of CAMSAP2 1214F and CAMSAP3(α-helix)-CAMSAP2 1239F. Data represent mean ± SEM. Asterisks represent significant differences as calculated by two-tailed Student’s t-test. n.s, P > 0.05; *P < 0.05; **P < 0.01; *** P < 0.001; **** P < 0.0001.

Therefore, we replaced the corresponding domains of CAMSAP1 and CAMSAP2 with the α-helix domain of CAMSAP3, we were surprised that the GFP signal, originally evenly distributed along the microtubules, became decorated at microtubule minus ends similar to the full length of CAMSAP3 or GFP-CAMSAP3 980F (Figure 1 D, 2 C and Figure 6 C-E). In contrast, when we replaced the α-helix domain of CAMSAP3 with the corresponding regions of CAMSAP1 and CAMSAP2, the corresponding minus ends decoration surprisingly disappeared, and a phenotype similar to the distribution pattern of CAMSAP1 or CAMSAP2 was obtained (Supplemental Figure 9). It should be pointed out here that since the MBD domain can also contribute to CAMSAP’s recognition of the minus end of microtubules (Supplemental Figure 7 A) (16), we used the C-terminus of the CAMSAP family to conduct the above experiments. These findings indicate that α-helix characteristics are required for CAMSAP3 to decorate at the minus ends of microtubules.

Finally, to have a more comprehensive understanding of the relationships between the MBD domains or α-helices of CAMSAP family proteins, we utilized phylogenetic analysis of CAMSAP/Patronin/PTRN-1 proteins across different species to reveal interesting evolutionary dynamics (34–36). Three distinct trees representing full-length proteins, α-helical domains, and the MBD domain revealed conserved and specialized features within the CAMSAP family. The full-length and MBD domain trees show strong conservation trends, particularly in CAMSAP2 and CAMSAP3, indicating essential, conserved functions in mammals, birds, and reptiles. In contrast, α-helical domain trees indicate unique evolutionary pathways within chordates, suggesting specialized roles distinct from the full-length protein lineage. This evolutionary snapshot highlights the functional significance of the CAMSAP domain, its structural conservation, and the adaptive specialization of protein fragments in response to cellular structural and dynamic demands (Supplemental Figure 10).

## Discussion

The CAMSAP family is comprised of non-centrosomal microtubule minus end binding proteins that play an important regulatory role in the microtubule network (2). Nezha/CAMSAP3 was the first identified CAMSAP family protein, which plays a crucial role in maintaining microtubule integrity by preventing depolymerization at minus ends (18, 19). Although CAMSAP family proteins can protect the minus ends of microtubules from depolymerization caused by kinesin13, the molecular mechanism by which CAMSAP itself dynamically regulates the minus ends of microtubules has not yet been made clear.

Recent advancements in cryo-electron microscopy have enabled milestone studies, such as Atherton et al. research on the key structural domain CKK of CAMSAP, which unveiled the molecular mechanism of microtubule minus end recognition from a structural biology perspective (22). Similarly, Liu et al. found that the C-terminus’s role in microtubule recognition and alteration of microtubule conformation marked a significant advancement in understanding the regulatory mechanisms of the CAMSAP family (26).

Our research, building on these foundational studies, delves into the dynamic regulatory molecular mechanism of the CAMSAP family, particularly focusing on microtubule minus ends. We have identified that the C-terminal α-helix of CAMSAP family proteins, especially in CAMSAP3, is critical for its function. This structural domain plays a key role in mediating the dimerization of the CKK domain, which not only enhances its microtubule-binding capability but also endows it with the ability to regulate microtubule dynamics. Additionally, the α-helix structure facilitates the specific distribution of CAMSAP at microtubule minus ends, thereby specifically regulating the dynamics of these ends.

### Dimerization Confers the Ability of CKK to Regulate Microtubule Dynamics

How does the CAMSAP family directly regulate microtubule dynamics? This question has long had no clear answer. As our previous reports and other researchers have revealed, the CKK domain is central to the form and function of CAMSAP proteins (22–24, 37) However, the CKK domain alone is unable to regulate microtubule minus end dynamics like full-length CAMSAP3, suggesting that additional factors are involved in this regulatory process (15, 16, 22, 37). We found that the α-helical domain of CAMSAP3 can mediate dimerization of CKK domain, thereby enhancing its affinity for microtubule binding (Figure 3). More importantly, this endows the CKK domain with the ability to regulate microtubule dynamics (Figure 5). However, we still do not understand its structural basis because the C-terminus of CAMSAP family proteins is rich in flexible regions and is therefore very difficult to observe with cryo-electron microscopy. We speculate that the ability of CKK to change microtubule conformation may play a crucial role in this process.

Furthermore, Atherton et al. have demonstrated that this related domain (976^th^ to the 1025^th^ a.a.) can interact with GMPCPP-stabilized microtubules (22). Similarly, Liu et al. reported that the D2 domain (975^th^ to the 1001^st^ a.a.) can bind to microtubules, influencing their conformation and thereby contributing to microtubule stability (26). Additionally, the MBD domain of CAMSAP3 could associate with the GMPCPP-stabilized seeds. Beyond direct regulation, CAMSAP3 collaborates with MCAK and KIF2A in modulating the dynamics of microtubule minus ends (13, 19). Thus, CAMSAP3’s influence on microtubule minus ends involves a complex interplay of multiple factors, reflecting the intricate regulation characteristic of CAMSAP family proteins. The potential role of CKK dimerization in these processes remains an intriguing question, which we aim to explore in future research.

### The α-helix Regulates CAMSAPs Decorate Microtubule Minus ends

The CKK domain is crucial for the recognition of CAMSAPs at the minus ends of microtubules (16, 22–24). When we initially reported Nezha (CAMSAP3), we found that the sequence at the C-terminus of CAMSAP3 impacts its localization at the minus end of microtubules (18). Meanwhile, Atherton et al. and Liu et al. reported that amino acids 976 to 1025 and 974 to 1111 of CAMSAP3, respectively, can directly bind to microtubules in vitro (22, 26). Notably, these sequences include the α-helix domain, underscoring that this region not only facilitates dimer formation but also plays a vital role in CAMSAP3’s ability to decorate microtubules. Moreover, our data indicate that the domain adjacent to the α-helix is crucial in determining the distinctive localization of CAMSAP family members at the minus ends of microtubules (Supplemental Figures 7 and Figure 6). By substituting the α-helical domain of CAMSAP1 and CAMSAP2 with that of CAMSAP3, the modified CAMSAP1 and CAMSAP2 exhibited intracellular localizations like CAMSAP3. This insight enhances our understanding of the variability in the decoration of CAMSAP family proteins at the minus ends of microtubules.

Furthermore, our results and previous reports also suggest that the MBD domain plays a critical role in this process (Supplemental Figure 7 B) (16, 22). It is very interesting that the MBD is relatively conserved in CAMSAP2 and CAMSAP3, which may also be a key factor in the ability of CAMSAP2 and CAMSAP3 to decorate microtubule minus ends very well, and the ability to accumulate here is very different from CAMSAP1. It’s worth noting that while both CAMSAP2 and CAMSAP3 have α-helical domains, the C-terminus of CAMSAP2 does not decorate the microtubule minus ends like CAMSAP3. Instead, it exhibits a distributed phenotype along microtubules (Figure 6 A and E). This indicates that the α-helix of CAMSAP3 possesses some unique properties (Figure 3 C). The decoration of microtubule minus ends is not closely related to its function in mediating dimer formation.

### Evolutionary Adaptation of the CAMSAP3 α-Helical Domain Pivotal Role in Microtubule Dynamics and Cellular Integrity

The CAMSAP family plays a pivotal role in the organization of non-centrosomal microtubules, significantly impacting cell structure and function (2). These proteins are indispensable in maintaining microtubule minus end dynamics, particularly in key areas such as neuronal differentiation, axonal regeneration, and cilia assembly, highlighting their importance across various biological contexts (37–41).

In vertebrates, the CAMSAP family comprises three members: CAMSAP1, CAMSAP2, and CAMSAP3, each displaying unique characteristics in their interactions and functions with microtubules (2). CAMSAP1 primarily tracks the microtubule minus ends, while CAMSAP2 goes beyond tracking to continue binding and decorating the extending minus ends, thus stabilizing the microtubules. CAMSAP2 and CAMSAP3 are distinguished by their highly conserved MBD domains and C-terminal regions, unlike CAMSAP1 (16, 22, 23). The presence of distinct protein domains (CKK, MBD, and α-helix) elucidates the varied behaviors of CAMSAP proteins at microtubule minus ends. CAMSAP3 demonstrates a more pronounced effect in regulating the dynamics of microtubule minus ends, especially due to its α-helical domain, which significantly enhances its binding affinity to microtubules, stabilizing and protecting the minus ends more effectively.

In conclusion, the α-helical domain of CAMSAP3 is an evolutionary adaptation to meet specific functional demands in microtubule dynamics, especially critical in cells dominated by non-centrosomal microtubules, such as epithelial cells. The dimerization tendency of the α-helix not only increases its affinity for binding to microtubules but also supports the cell’s vitality and functionality by stabilizing microtubules. The conservation of this domain across various vertebrates underscores its importance in evolution and its key contribution to maintaining cell integrity and functionality. In summary, the α-helical domain of CAMSAP3 represents a significant evolutionary innovation in response to the needs of cell structure and functionality, reflecting its central role in the regulation of microtubule dynamics.

## Materials and Methods

The DNA constructs, antibodies, and reagents used in this study are described in SI Appendix, Materials and Methods. The detailed procedures of cell culture and transfection, immunofluorescence staining, living-cell imaging, Co-IP, protein expression and purification, in vitro microtubule dynamics assay, in vitro microtubule co-sedimentation assay, SEC-MALS, structure prediction, protein sequence alignment and evolutionary analysis, data and statistical analysis are provided in Appendix, Materials and Methods.

## Acknowledgments

We thank Masatoshi Takeichi for his advice. We thank members of the Meng group for helpful discussions. We thank Xing Jia, Qing Bian, Xiang Zhang, and Yun Feng for technical support with Confocal imaging, Multi-SIM images, and image analysis. We thank Shan Sun and Huabing Ruan for advice about structure analysis. This work was funded by the National Natural Science Foundation of China (31930025 and 32070704), the National Key Research and Development Program of China (2021YFA0804802) and the National Natural Science Foundation of China (32100760).

## Author Contributions

Y.L., R.Z., X.L., and W.M. designed all experiments, interpreted the results, and prepared the manuscript. J.R. and D.L. performed the static light detection and analyzed the predicted structure. Y.L. and R.Z performed microtubule dynamics assays with the help of W.C. and H.C. Z.Z. purified tubulin protein from the porcine brain. H.X. contributed to technical support. Q.X., W.J., and W.F. provided reagents and advice.

